# Insulin modulates mPFC gene expression and emotional behavior in a sex-specific manner following fetal growth restriction

**DOI:** 10.1101/2025.08.14.669951

**Authors:** Patrícia Maidana Miguel, Barbara Barth, Aashita Batra, Márcio Bonesso Alves, Danusa Mar Arcego, Bonnie Alberry, Roberta Dalle Molle, Ameyalli Gómez-Ilescas, Tie Yuan Zhang, Xianglan Wen, Carine Parent, Nicholas O’Toole, André Krumel Portella, Patrícia Pelufo Silveira

## Abstract

Exposure to prenatal adversity (e.g., stress, malnutrition) is a major risk factor for lifelong vulnerability to neuropsychiatric and metabolic disorders, and alterations in the function of peripheral hormones in the brain are suggested as a possible and unexplored mechanism. Although insulin is well known for its peripheral metabolic functions, it also influences brain development and emotional regulation, particularly within the medial prefrontal cortex (mPFC). Here, we identify insulin signaling as a key mechanistic link between prenatal stress and long-term alterations in brain function and behavior. Using a validated rat model of prenatal adversity (prenatal food restriction, FR), we found that insulin administration selectively modulates gene expression in FR animals at P0, P21 and P90, affecting pathways involved in neurodevelopment and stress regulation, such as Wnt/β-catenin signaling. Targeted insulin infusion into the mPFC of adult animals reversed behavioral phenotypes induced by FR in a sex- and context-dependent manner, decreasing emotional reactivity to environmental cues. These findings offer novel insight into the neurodevelopmental role of insulin and suggest that insulin signaling may serve as a therapeutic target to mitigate the long-term brain and behavioral consequences of prenatal adversity.

## Introduction

Globally, one in four individuals experiences significant intrauterine adversities that impact their growth and development ^1^, as well as their risk for both psychiatric and metabolic conditions later in life ^2^. For instance, individuals born small for gestational age exhibit a heightened risk of developing anxiety disorders in adulthood ^3–5^. Fetal growth restriction (FR) also impairs pancreatic β-cell development ^6^, leading to basal dysregulation of insulin secretion and reduced glucose responsiveness ^7^. This dysregulation contributes to altered insulin sensitivity, glucose intolerance, and elevated risk for type 2 diabetes ^8, 9^.

Beyond its peripheral metabolic functions, insulin signaling in the brain influences mood and cognitive processes ^10–12^. Insulin crosses the blood-brain barrier via active transport and provides the brain with information about the status of long-term energy stores ^13^, while also modulating neurotransmission ^14, 15^. Although insulin receptors are broadly expressed across the brain, their role in regulating emotional behavior is especially relevant in the medial prefrontal cortex (mPFC). In rodent models of diabetes, mPFC neuronal loss, apoptosis, and volume reductions have been associated with increased anxiety-like behavior ^16,17^. In individuals with type 2 diabetes, metformin (an antidiabetic drug) treatment induces a lower incidence of anxiety disorders ^18, 19^.

Supporting a mechanistic link, metformin treatment activates AMP-activated protein kinase (AMPK) in the mPFC and mitigates social stress-induced anxiety-like behaviors in rodents ^20^.

Insulin has been proposed as a key mediator linking prenatal adversity to neurodevelopmental alterations and long-term brain function. Disruptions in insulin signaling may underlie the association between early adversity and increased risk for psychiatric and cardiometabolic conditions ^21, 22^. Insulin resistance has been consistently observed in individuals born small for gestational age ^8, 9^, as well as in rodent models of fetal growth restriction ^23–25^. In humans born small, insulin resistance is associated with altered behavioral responses to environmental stimuli, including heightened psychological distress, anxiety/depressive and social problems ^26–28^, lower tolerance for delayed gratification and impairments in value-based decision-making ^29–31^. FR in humans is also linked to altered hedonic processing, such as blunted reactivity to sucrose, increased food intake, and elevated PFC activation in response to food-related cues ^32–34^, effects that are modulated by peripheral insulin resistance ^32^.

These alterations in behavioral and emotional responses to environmental stimuli may contribute to long-term vulnerabilities in individuals born small, affecting daily choices and overall health. Similarly, FR rodents show a preference for smaller, immediate rewards over larger, delayed ones ^35^, increased anxiety-like behaviors ^36^, and altered hedonic and conditioned responses to sweet foods ^37, 38^. These animals exhibit reduced dopamine release in the nucleus accumbens and PFC when exposed to palatable foods ^24, 35^, an effect that is reversed by peripheral insulin administration ^24^. These findings align with evidence that dopamine neurons are responsive to novel or salient stimuli ^39^, are shaped by early adversity ^40^, and are modulated by central insulin signaling (review in ^22^).

The associations between FR and altered behavioral responses to the environment are strongly influenced by mPFC neurotransmission and modulated by insulin, with significant implications for health across the lifespan. It is therefore essential to determine whether altered insulin function in FR individuals specifically affects the mPFC transcriptional landscape and contributes to changes in emotional reactivity and anxiety-like behaviors. Here, we examined how peripheral insulin modulates mPFC gene expression profiles of FR animals at different developmental stages: adults (P90), juveniles (P21), and neonates (P0). We then confirmed that mPFC insulin administration affects behavioral responses to the environment in tasks that evaluate emotionality in a new setting (spontaneous alternation and light-dark box tests). We hypothesize that insulin acts as a key modulator of mPFC development and function, shaping emotional/anxiety-like behaviors, particularly in animals exposed to fetal growth restriction.

## Materials and methods

Sprague Dawley rats (Charles River Laboratories Inc, St. Constant, QC) were exposed to prenatal food restriction (FR) or *ad libitum* feeding from gestational day 10 to birth, after which all litters were fostered to *ad libitum*-fed dams. The offspring were randomly injected with either saline (1mL/kg) or insulin (5IU/mL in saline) by intraperitoneal injection (i.p.), and had their brain collected after 30 minutes, in three distinct developmental timepoints: postnatal day (P) 0, 21, and 90. The medial prefrontal cortex (mPFC) was isolated by tissue punch from fresh frozen coronal brain sections for bulk RNA sequencing (n = 9-10/group/sex/timepoint; GEO accession GSE295966). Another offspring cohort received a bilateral 1-μl solution of insulin (0.25 μg/μl) or a 0.9% sterile saline solution at the mPFC in adulthood (after P90), and 5 minutes later were exposed to the spontaneous alternation (SAT) and light-dark box (LDB) tests. Behavioural performance was analyzed using Generalized Linear Model (GLM) and a Linear Mixed Model (LMM). Methods are fully detailed in the Supplement and outlined in Fig. 1A, 5A, Supplemental Fig. 2A and 3A. All procedures comply with the ethical standards of the national guide on the care and use of laboratory animals and have been approved by the Douglas Mental Health University Institute – AUP DOUG-8018.

**Fig. 1:**
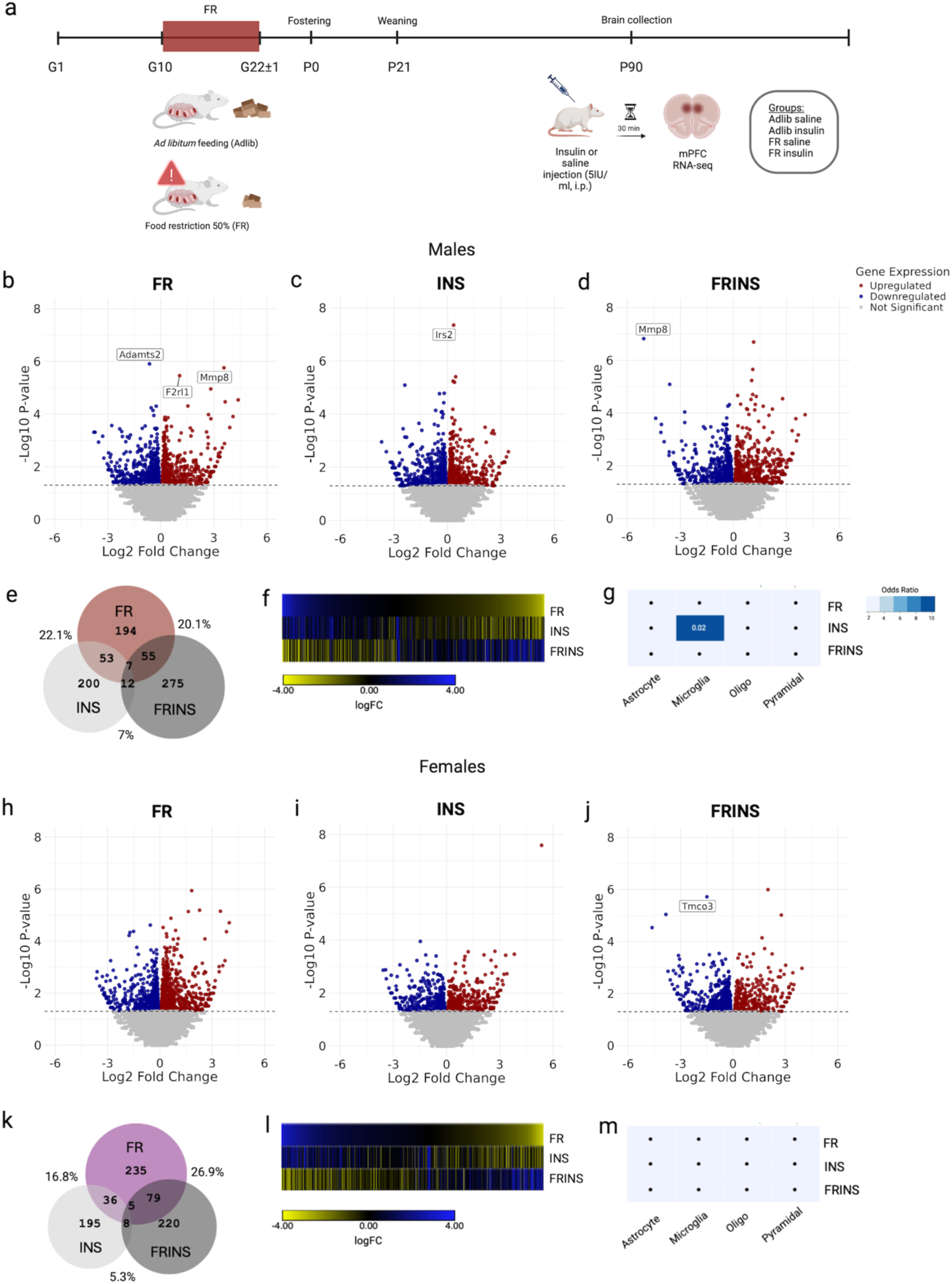
Insulin injection induces mPFC transcriptional changes dependent on the prenatal adversity condition and sex. Schematic timeline of experimental procedures, including the prenatal adversity model of food restriction (FR) from gestational day 10 (G10) until the end of gestation, and offspring mPFC collection for RNA-sequencing analysis in postnatal day 90 (P90) **a**. Volcano plots showing differentially expressed genes (DEGs) (FDR < 0.05) associated with FR, INS, and FRINS conditions in males **b-d** and females **h-j.** Venn diagrams represent the number of DEGs (uncorrected p < 0.05, fold change > 30%, and protein-coding genes) altered by fetal-growth restriction (FR), insulin injection in control animals (INS), or insulin injection in FR animals (FRINS). Percentage values represent the extent of overlap between conditions **e,k**. Union heatmaps display the log fold change of DEGs across matched comparisons, irrespective of statistical significance. Yellow indicates increased gene expression (positive log fold change), while blue denotes decreased expression (negative log fold change) **f,l**. Cell-type enrichment analysis of DEGs in each comparison using a curated list of cell-type-specific genes expressed in the PFC **g,m.** G = gestational day; PND = postnatal day. DEGs analysis considers the following comparisons: FR = saline injection in fetal growth restriction vs saline injection in adlib animals, INS = insulin injection in adlib animals vs. saline injection in adlib animals, FRINS = insulin injection in FR animals vs saline injection in FR animals. n = 9-10/group/sex.

## Results

### Prenatal growth restriction induces sex-specific changes in mPFC gene expression related to environmental responsivity

We first investigated how the metabolic state under basal conditions or following peripheral insulin administration shapes transcriptional responses in the adult mPFC of FR animals (P90) (Fig.1a). We identified differentially expressed genes (DEGs) associated with FR (FR vs. Adlib) under saline conditions, analyzed separately by sex. Using false discovery rate (FDR) correction < 0.05, we found that in FR males, Mmp8 and F2rl1 were upregulated, while Adamts2 was downregulated (Fig.1b). In contrast, no significant DEGs were identified in FR females (Fig.1h). These results suggest that FR induces male-specific transcriptional alterations in mPFC genes related to stress susceptibility and metabolic dysregulation, highlighting sex-specific mPFC molecular signatures of prenatal adversity.

### mPFC insulin sensitivity is altered in prenatally growth-restricted males

To investigate insulin sensitivity in the mPFC, we identified DEGs associated with insulin injection (insulin vs. saline) in both Adlib and FR animals. In Adlib males, insulin injection induced upregulation of Irs2 (Insulin receptor substrate 2) (Fig.1c), but this effect was absent in FR males (Fig.1d), suggesting reduced responsiveness to insulin following fetal growth restriction. Interestingly, in FR males, insulin downregulated Mmp8 – which was upregulated in FR saline – reversing the effects of prenatal adversity on Mmp8 expression in males (Fig.1d). In FR females, insulin induced downregulation of Tmco3 (also known as Tmcc3) (Fig.1j). FR is linked to differential gene expression in genes related to emotional responsivity to environmental stress, insulin resistance and neuroinflammation, while insulin administration reverts these patterns.

### Partial overlap in the mPFC transcriptional signature between insulin administration and prenatal growth restriction

To examine broader transcriptional patterns in the mPFC, we identified DEGs with a nominal *p* < 0.05 and a log₂(fold change) > |0.3785| (corresponding to a fold change > 30%) ^41^. In males, there was a 22.1% overlap between DEGs induced by insulin injection (vs. saline in Adlib animals) and those altered by fetal growth restriction (FR vs Adlib under saline conditions) (Fig.1e); in females, this overlap was 16% (Fig.1k). Union heatmaps sorted by FR-regulated genes further demonstrated that FR and insulin induced transcriptional changes in the same direction in both males (Fig.1f) and females (Fig.1l). In males, insulin-regulated genes in Adlib were significantly enriched in microglia (Fig.1g), while no significant enrichment was found in females (Fig.1m) or in FR-regulated genes in either sex. Insulin administration mimics some of the transcriptional changes induced by FR in the mPFC, supporting the hypothesis of a latent insulin resistance state following FR.

### Insulin and saline have opposite effects on mPFC transcriptional signatures in prenatally growth-restricted animals

We examined how insulin administration alters gene expression in the mPFC of FR animals by comparing insulin-treated FR animals (FRINS) to their saline-treated FR counterparts. There was a 20% DEG overlap in males (Fig.1e) and 26% in females (Fig.1k). Union heatmaps showed inverse patterns of gene expression between the conditions in both sexes (Fig. 1f,l), suggesting that FR animals exhibit chronic, insulin-resistant adaptations at baseline, which are reversed by an acute, supraphysiologic insulin challenge. No significant cell population enrichment was found in the FRINS condition in either sex (Fig.1g,m). Insulin-induced transcriptional changes in Adlib and FR animals shared minimal overlap, with only 5– 7% of DEGs in common across both sexes (Fig.1e,k), which was confirmed by union heatmaps (Fig.1f,l). These findings indicate that insulin exerts distinct and largely non-overlapping transcriptional effects in the mPFC of FR animals compared to controls.

### WNT/β-catenin signaling in embryogenesis is a common pathway in response to FR, INS, and FRINS in females, and to FR in males

We sought to identify unique or shared pathway maps across the three conditions (FR, INS, FRINS) within each sex. In males, the WNT/β-catenin signaling in embryogenesis pathway emerged as uniquely associated with FR (Fig.2a). In females, WNT/β-catenin signaling was also uniquely associated with FR, but notably, it was the only pathway shared across FR, INS, and FRINS (through alterations in Wnt ligands), underscoring its relevance to both prenatal adversity and insulin sensitivity effects (Fig.2b,c). Uniquely in females, FR was linked to pathways involved in lipid and lipoprotein synthesis and metabolism (Fig.2b), whereas in males, these pathways were uniquely associated with insulin (Fig.2a). Together, these results highlight WNT/β-Catenin signaling as a core mPFC pathway responsive to prenatal adversity in both sexes, and a convergent pathway across FR, INS, and FRINS in females (Fig.2c).

**Fig. 2:**
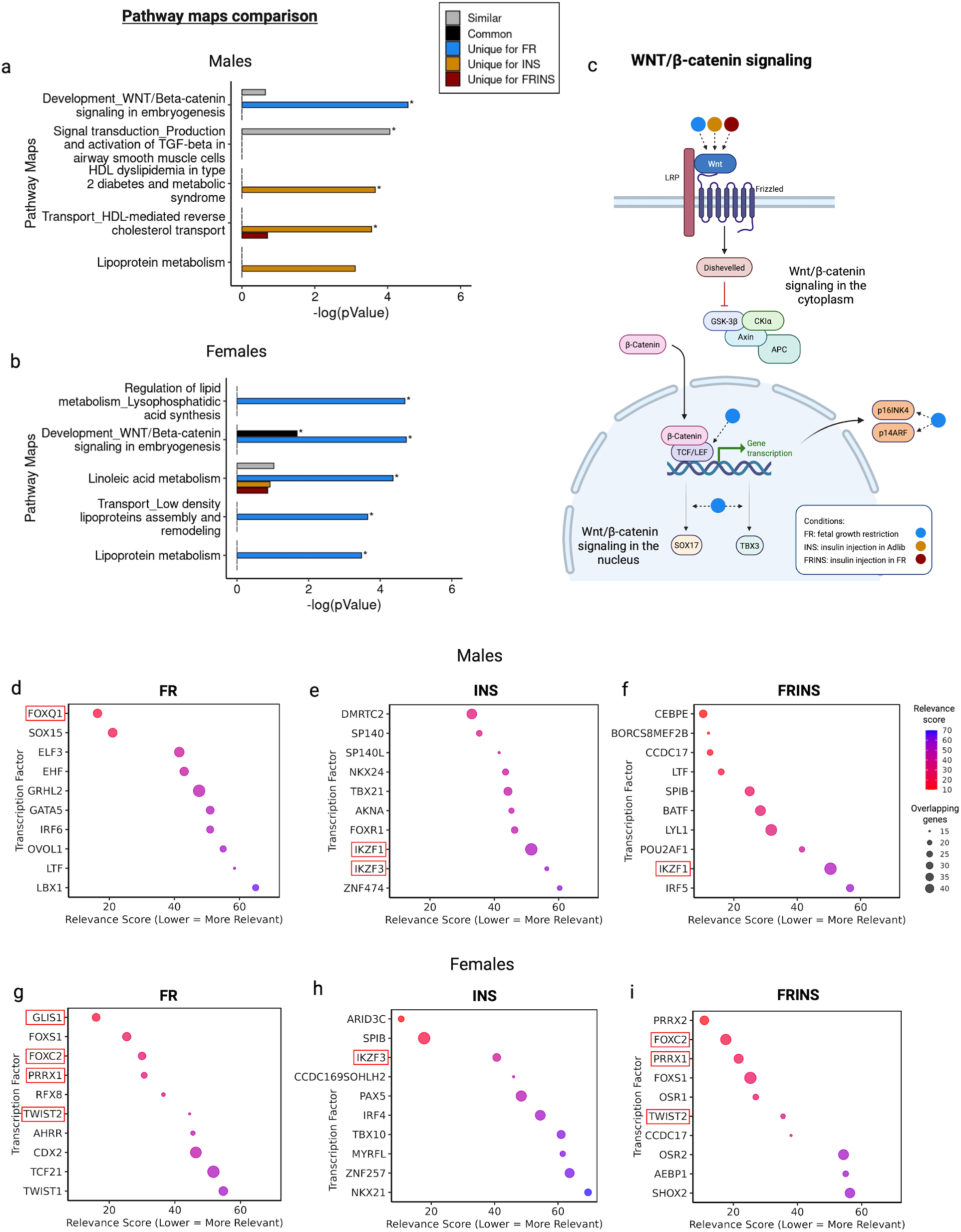
Pathwap map analyses revealed WNT/β-catenin signaling in embryogenesis as a common pathway responsive to FR, INS, and FRINS conditions in females. Pathway map analyses identified unique, similar, and common pathways associated with the DEGs of each condition, analyzed separately in males and females. WNT/β-catenin signaling in embryogenesis is uniquely affected by fetal growth restriction in both males and females, and in females, all three experimental conditions (FR, INS, and FRINS) affect the expression of Wnt ligands **a,b.** Canonical WNT/Beta-catenin signaling pathway plays key roles in animal development. WNT proteins bind to Frizzled receptors and their co-receptors LRP, leading to the activation of Dishevelled. This activation inhibits the cytoplasmic “destruction complex”, composed of glycogen synthase kinase-3β (GSK-3β), axin, adenomatous polyposis coli (APC), and casein kinase I (CK1), which degrade β-catenin. Inhibition of this complex stabilizes β-catenin, allowing it to translocate to the nucleus, where it activates gene expression through TCF/LEF transcription factors. In our findings, all three experimental conditions (FR, INS, and FRINS) affect the initiation of the Wnt/β-catenin cascade in females, though through different genes: Wnt5a in FR, Wnt9a in INS, and Wnt2b and Wnt9b in FRINS. Fetal growth restriction, however, also modulates downstream components of the pathway, such as Lef-1, Tcf(Lef), TBX3, SOX17, p16INK4, and p14ARF **c.** Top ten transcription factors (TFs) associated with the DEGs of each condition are ranked by relevance score, where lower means higher relevance, and bubble sizes represent the number of overlapping target genes per TF **d-i.** DEGs analysis considers the following comparisons: FR = saline injection in fetal growth restriction vs saline injection in adlib animals, INS = insulin injection in adlib animals vs. saline injection in adlib animals, FRINS = insulin injection in FR animals vs saline injection in FR animals.

### Prenatal growth restriction-associated mPFC gene expression is linked to transcription factors regulating fetal growth, type 2 diabetes, neuroinflammation, and stress responses

We observed a distinct pattern of TFs associated with DEGs across conditions and sex. FOXQ1, known to modulate neuroinflammation and influence cognitive performance ^42^ appears as the most relevant TF in FR males (Fig.2d). In FR females, GLIS1 emerged as the top-ranked TF (Fig.2f). The GLIS family (GLIS1–3) has been implicated in intrauterine growth restriction, pancreas development, neurodegenerative disease, and type 2 diabetes ^43^. Members of the Ikaros family (IKZF1 and IKZF3), suggested as a link between stress and immune responses ^44^ were associated with the DEGs induced by insulin in both control and FR males and females (Fig.2e,g,h). In FR females (both FR and FRINS), FOXC2 – a key regulator of adipocyte metabolism, insulin resistance, and gestational diabetes ^45^ – was consistently identified as a top TF (Fig.2g,i). Other TFs strongly associated with FR and FRINS in females included TWIST2, involved in TNF-α regulation ^46^, and PRRX1, which is required for self-renewal and proper differentiation of cortical neural progenitors ^47^ (Fig.2g,i).

### Developmental trajectories of mPFC transcriptomic signatures in response to FR and insulin

We subsequently performed RNA-sequencing at two key time points: after the catch-up growth period (P21) and at birth (P0), immediately following the stress exposure (Sup. Figs.2a,3a). The total number of DEGs was lower at P0 and P21 compared to P90 (Fig.3a-c). Two-sided rank–rank hypergeometric overlap analysis (RRHO) ^48^ was used to assess patterns and strengths of genome-wide overlap between the groups, sex and ages. The analyses at P90 confirm that there is a similar pattern of gene regulation by FR and insulin administration in both males (Fig.3d, left) and females (Fig.3e, left), with stronger concordance in males. At P90, insulin exerts a regulation of gene expression in the opposite direction of saline in FR animals in both sexes (Fig. 3d,e; middle), with a stronger discordant pattern in females. The response induced by insulin administration on mPFC gene expression in FR animals is completely unrelated to the effect induced in control animals (Fig.3d,e; right).

**Fig. 3:**
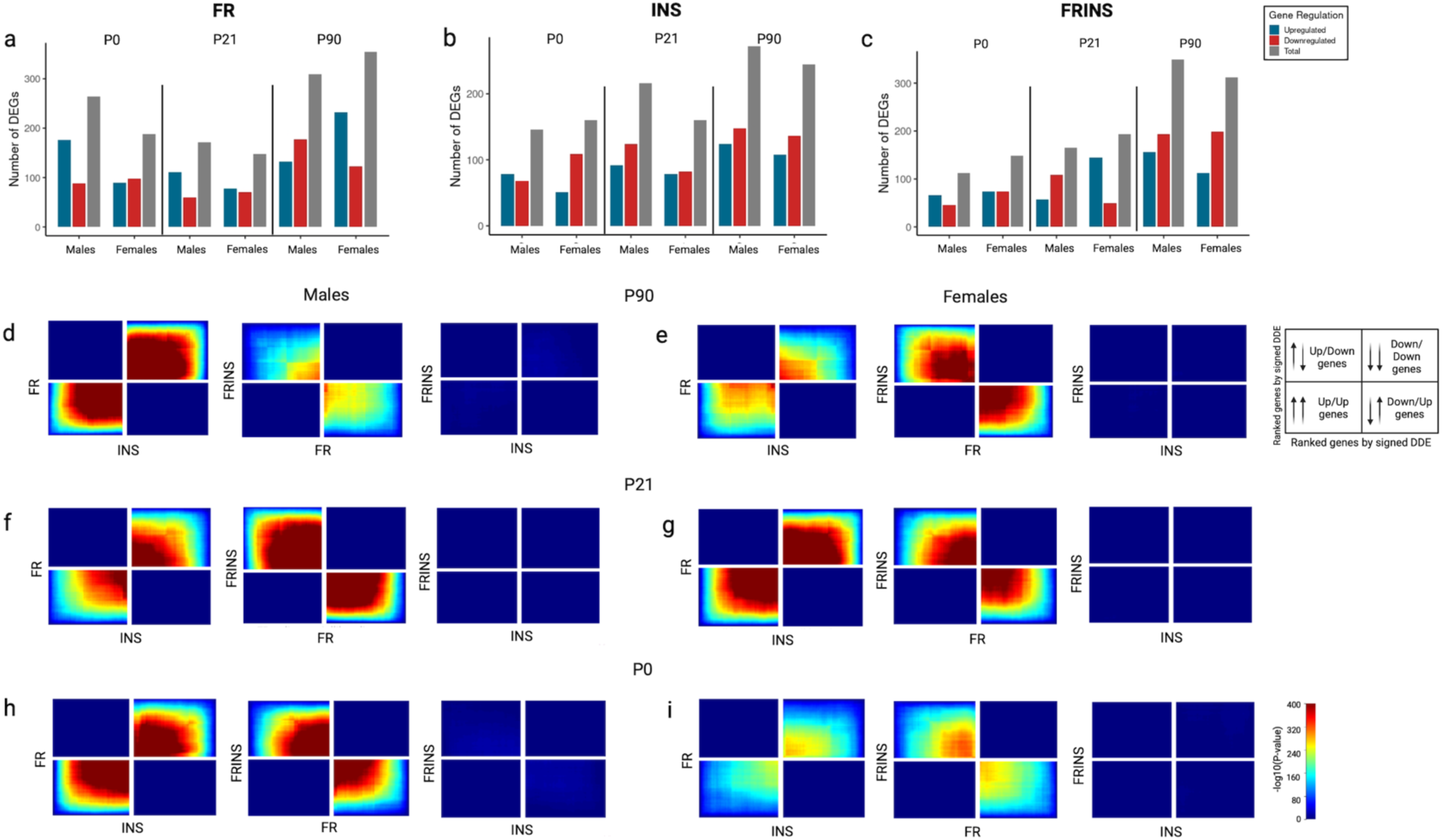
Developmental trajectories of mPFC transcriptomic signatures in response to prenatal growth restriction and insulin. Bar plots representing the number of upregulated, downregulated, and total number of DEGs in males and females for each timepoint (P0, P21, P90) in the three distinct conditions (FR, INS, and FRINS) **a-c**. Threshold-free comparison of DEGs using rank-rank hypergeometric overlap (RRHO) analysis in the distinct timepoints **d-i.** Each pixel represents the significance of overlap (−log₁₀ p-value from a hypergeometric test) between the transcriptomes of the indicated comparisons. Color intensity reflects overlap strength. Co-upregulated genes appear in the lower left quadrant, co-downregulated genes in the upper right, and oppositely regulated genes in the upper left (up–down) and lower right (down–up) quadrants. Genes along each axis are ranked from most to least significantly regulated, progressing from the center outward. DEGs analysis considers the following comparisons: FR = saline injection in fetal growth restriction vs saline injection in adlib animals, INS = insulin injection in adlib animals vs. saline injection in adlib animals, FRINS = insulin injection in FR animals vs saline injection in FR animals.

RRHO analyses at P21 and P0 closely mirrored the P90 patterns (Fig.3f-i). The degree of agreement induced by FR and by insulin administration is somewhat stable in males during development (Fig.3d,f,h; left), while in females, it is very mild at P0, stronger at P21 and moderate at P90 (Fig.3e,g,i; left). When comparing the effect of insulin vs. saline in FR during development, opposite regulation is present in males at P0 and becomes much stronger at P21, declining in adulthood (Fig.3d,f,h; middle); in females, there is a very discreet opposite regulation at P0, which gets more evident and stable at P21 and P90 (Fig.3e,g,i; middle).

When examining overlap in upregulated and downregulated DEGs across development (P0, P21, and P90), a few DEGs showed consistent directionality across two timepoints, but no DEG was consistently altered across all three stages (Sup. Fig.1) — highlighting the dynamic nature of mPFC gene expression throughout postnatal development. Compared to the P90 analyses, the proportion of overlapping DEGs between FR and INS, as well as between FR and FRINS, remained similar at earlier timepoints (Sup. Fig.2e,k; Sup. Fig.3e, k). Notably, the minimal DEG overlap between INS and FRINS observed at P90 was consistently maintained at earlier time points, with a maximum overlap of only 8.5% in P21 males (Sup. Figs.2e,k and 3e,k), reinforcing the idea that insulin has markedly different effects on mPFC gene expression in FR animals compared to Adlib.

### Insulin administration upregulates genes related to thyroid function specifically in FR females at earlier ages

A more detailed analysis of FDR-corrected DEGs at earlier developmental stages revealed a downregulation of Cyp2g1 and Cyp2a2, members of the cytochrome P450 (Cyp) 2 family, in response to insulin in males at P21 (Sup. Fig.2d). In contrast, FR females showed upregulation of these same genes in response to insulin (Sup. Fig.2j). Importantly, FR in females was associated with the downregulation of Car3 (Sup. Fig.2h), whereas insulin response in FR related to the upregulation of this gene (Sup. Fig.2j), suggesting a reversal of this prenatal adversity-induced effect. FR in females was also associated with downregulation of Fam111a, which encodes a protease involved in DNA replication, transcription, and microtubule integrity ^49^ (Sup. Fig.2h). Insulin treatment in FR females led to the selective upregulation of genes involved in thyroid hormone signaling, including Trh (thyrotropin-releasing hormone) and Ttr (transthyretin, a thyroid hormone transport protein) (Sup. Fig.2j). Insulin injection in FR also upregulated genes related to stress regulation, such as S100a5 (a calcium-binding protein involved in stress and immune responses ^50^), Bpifb3 (a lipid-binding protein), and Eomes (a transcription factor essential for neurogenesis ^51^) (Sup. Fig.2j). At P0, the only significant DEG was the upregulation of Gh1 (growth hormone 1) in response to insulin in females (Sup. Fig.3i), further suggesting that insulin may stimulate endocrine-related gene expression early in development.

### mPFC gene expression induced by insulin is linked to neuroendocrine peptides, oligodendrocyte differentiation, and immune response at early developmental stages

We next examined common and unique biological pathways associated with mPFC transcriptional signatures during early development (Fig.4). At P21 in males, FR was significantly associated with pathways related to pro-opiomelanocortin (POMC) processing and function (Fig.4a). In contrast, acute insulin treatment at this stage significantly impacted pathways related to oligodendrocyte differentiation and type 2 diabetes, suggesting early modulation of both neural and metabolic pathways (Fig.4a). In females at P21, none of the three conditions elicited statistically significant pathway enrichment (Fig.4b). Similarly, at P0, pathway enrichment was generally less robust (Fig.4i, j). Although not statistically significant, WNT/β-catenin signaling emerged as a shared pathway across FR, INS, and FRINS in females at P0 (Fig.4j), mirroring its consistent role observed in later developmental stages. The top transcription factor (TF) associated with FR in P21 males was Tbx21, which plays a key role in immune cell differentiation and function and has been implicated in insulin sensitivity^52^ (Fig.4c), and was also relevant in FRINS (Fig.4e). In females at P0, FOXJ1 was associated with genes altered across all three conditions (FR, INS, and FRINS) (Fig.4n-p). FOXJ1 has been previously shown to be dysregulated in the mPFC in models of palatable food intake and cocaine addiction ^53^, suggesting it may be involved in neuroplastic adaptations to environmental stimuli. Interferon regulatory factors (IRF4, IRF5, and IRF8) were identified as relevant TFs in FRINS males at both P21 and P0 (Fig.4e,m), as well as in INS males at P0 (Fig.4l), indicating a potential role in neuroimmune responses under both conditions. Finally, Hnf4a was a key TF in FRINS males and females at P0 (Fig.4m,p). Hnf4a is a well-known regulator of islet development, β-cell function, and glucose metabolism, including the regulation of genes involved in glucose transport and glycolysis ^54^.

**Fig. 4:**
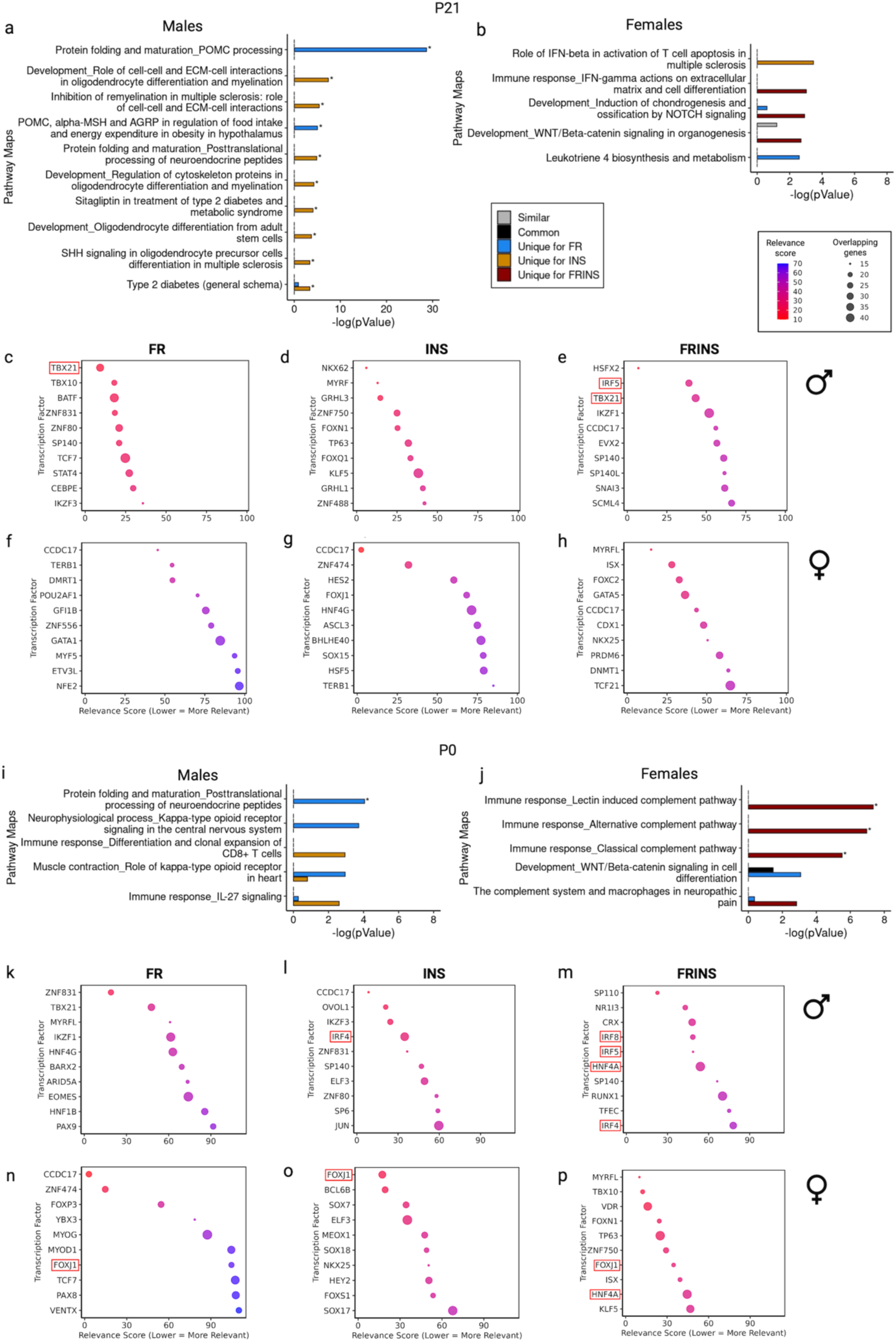
Pathway map analyses revealed key pathways associated with FR, INS, or FRINS in early developmental stages. Pathway map analyses identified unique, similar, and common pathways associated with the DEGs of each condition, analyzed separately in males and females in P21 **a,b** or P0 **i-j.** Top ten transcription factors (TFs) associated with the DEGs of each condition in P21 and P0 are ranked by relevance score, where lower means higher relevance, and bubble sizes represent the number of overlapping target genes per TF **c-h, k-p.** DEGs analysis considers the following comparisons: FR = saline injection in fetal growth restriction vs saline injection in adlib animals, INS = insulin injection in adlib animals vs. saline injection in adlib animals, FRINS = insulin injection in FR animals vs saline injection in FR animals.

### mPFC insulin-associated anxiolytic effect depends on prenatal growth restriction and sex

We then investigated whether insulin infusion in the mPFC modified the spontaneous alternation in FR and controls (Fig.5a). There was a main effect of sex, with females displaying a higher number of spontaneous alternation and total arm entries (Fig.5b,c) than males in the total 15-min performance. The spontaneous alternation percentage (relative to total entries) was not affected by any condition in the overall performance (Fig.5d). To better understand temporal dynamics and group-specific responses, we analyzed performance over time separately in males and females. In males, mPFC insulin injection increased both the number of spontaneous alternation (Fig.5e) and total entries (Fig.5f) over time, indicating enhanced exploratory behaviour, regardless of the prenatal experience. The number of spontaneous alternations over 15 minutes in FR males suggests a differential response to the new apparatus, although the comparison with controls does not reach statistical significance (time x adversity interaction, p=0.07) (Fig.5e). Among females, no significant group differences were observed over time (Fig.5h,i,j).

**Fig. 5:**
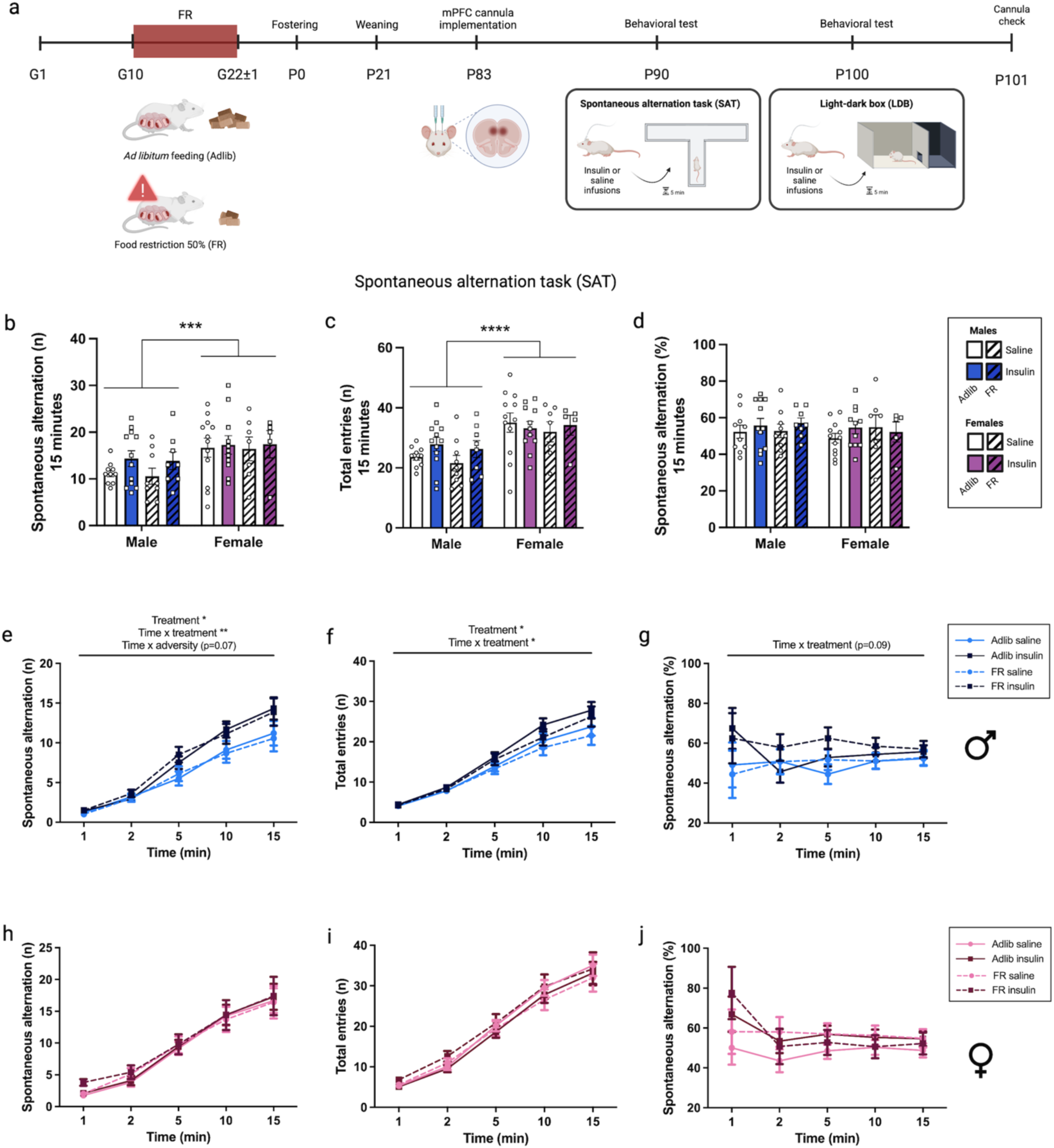
Insulin administration in the mPFC increases spontaneous alternation (SA) in males independent of the adversity condition. Schematic timeline of experimental procedures, including the prenatal adversity model of food restriction (FR) from gestational day 10 (G10) until the end of gestation, and offspring behavioral analyses in adulthood **a**. Total number of SA over the 15 minutes in the spontaneous alternation task (SAT) elicits a sex main effect (Wald χ²(1) = 11.08, p < 0.001) **b**. Total number of arm entries over the 15 minutes also shows sex effects, suggesting that females have more exploratory behavior than males (Wald χ²(1) = 22.54, p < 0.0001) **c**. Percentage of SA has no significant effect between groups **d.** Cumulative number of SA over the 15 minutes in the SAT in males showed a significant treatment effect (*F*_1, 37.14_ = 5.46, p = 0.025), and a time x treatment interaction effect (*F*_4, 55.19_ = 3.88, p = 0.008), but time x adversity interaction effect did not reach statistical significance (*F*_4, 55.19_ = 2.29, p = 0.071) **e**. Cumulative number of total entries also elicited treatment (*F*_1, 39.25_= 5.83, p = 0.021) and time x treatment interaction (*F*_4, 55.40_ = 3.50, p = 0.013) effects in males, with insulin increasing the exploratory behavior in both groups **f**. Cumulative percentage of SA over the 15 minutes in males, which is corrected by the total number of entries, had a non-statistically significant time x treatment interaction (*F*_4, 69.08_ = 2.10, p = 0.09) **g**. Cumulative number of SA, number of entries, and percentage of SA over the 15 minutes in the SAT in females demonstrated no significant group effects, except for a main effect of time across all measures (Entries: *F*_4, 46.04_ = 65.53, p < 0.0001; SA: *F*_4, 47.58_ = 64.73, p < 0.0001; SA%: *F*_4, 74.39_ = 3.09, p = 0.021), indicating an increase in SA and total entries, and a decrease in SA percentage **h, i, j**. A generalized linear model (GLM) was applied for overall 15-minute analyses, including both sexes, whereas a linear mixed model was applied to investigate the performance throughout time separately in males and females. * *p* < 0.05, ** *p* < 0.01, *** *p* < 0.001,**** *p* < 0.0001. G = gestational day; P = postnatal day. Data are expressed as mean±S.E.M; n = 5-12/group.

We next evaluated whether mPFC insulin infusion affected anxiety-like behavior using a 15-minute light-dark box test, which presents a conflict between the drive to explore and the aversion to brightly lit areas ^55^. There was a significant adversity x treatment interaction where insulin increased time spent on the light side in FR animals, compared to both controls and saline-treated animals, in both sexes (Fig.6a). This suggests that mPFC insulin infusion 5 minutes before the task has an anxiolytic effect that is specific to FR animals, regardless of sex. For the number of entries into the lit side (Fig.6b) and risk evaluation percentage (Fig.6c), only sex effects were found.

**Fig. 6:**
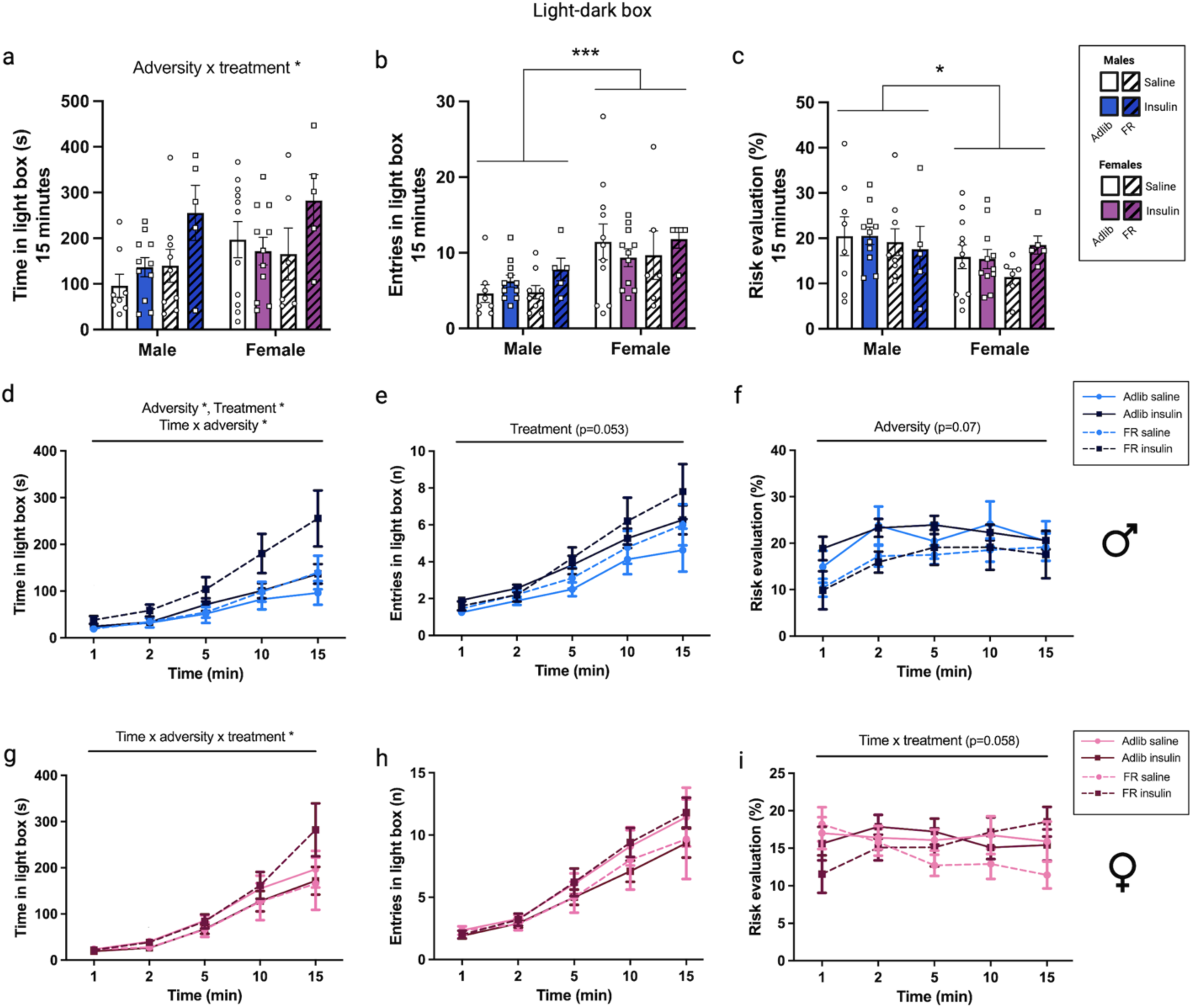
Insulin administration in the mPFC differently affects performance in the light-dark box (LDB) depending on the adversity group and biological sex. Total time in the light box over the 15 minutes in the LDB elicits an adversity x treatment interaction effect (Wald χ²(1) = 4.22, p = 0.040), indicating that insulin increases the time in the lit side in the adversity group **a**. Total number of entries in the light box elicited sex effects (Wald χ²(1) = 13.75, p < 0.0001) **b**, similar to the risk evaluation percentage (Wald χ²(1) = 4.12, p = 0.042) **c**, indicating that females entered more in the lit side and performed less risk evaluation. Cumulative time spent over the 15 minutes in the LDB in males showed adversity (*F*_1, 30.16_ = 4.35, p = 0.046), treatment (*F*_1, 30.16_ = 5.48, p = 0.026), and time x adversity (*F*_4, 49.95_ = 3.02, p = 0.026) interaction effects, suggesting that both FR and insulin increased the time in the light over time **d**. Insulin induced a non-significant increase number of entries over 15 minutes in the LDB in males (*F*_1, 31.58_ = 4.05, p = 0.053) **e**. Cumulative risk evaluation percentage over the 15 minutes in the LDB in males suggests that FR groups performed less risk evaluation, but this effect did not reach statistical significance (*F*_1, 27.58_ = 3.37, p = 0.077) **f**. Cumulative time spent over the 15 minutes in the LDB in females showed time x adversity x treatment interaction effect (*F*_4, 47.49_ = 3.68, p = 0.011), indicating that insulin treatment increased time in the light, especially in FR animals towards the end of the task **g**. Cumulative number of entries over the 15 minutes in the LDB in females had no significant group effects **h**. Cumulative risk evaluation percentage over the 15 minutes in the LDB in females showed a non-statistically significant time x treatment interaction (*F*_4, 84.9_ = 2.37, p = 0.058). **i**. A generalized linear model (GLM) was applied for overall 15-minute analyses, including both sexes, whereas a linear mixed model was applied to investigate the performance throughout time separately in males and females. * *p* < 0.05, ** *p* < 0.01, *** *p* < 0.001,**** *p* < 0.0001. G = gestational day; P = postnatal day. Data are expressed as mean±S.E.M; n = 5-12/group.

When behavior was analyzed over time, both adversity and insulin treatment increased time spent in the light compartment in males (Fig.6d), consistent with anxiolytic effects. Notably, FR insulin-treated males showed the greatest increase in the lit-side exploration (Fig.6d). In females, a time × adversity × treatment interaction effect was observed for time spent in the light compartment (*p* = 0.011; Fig.6g). FR-saline females have lower average time spent in the lit side in comparison to Adlib-saline females, and this pattern is completely reversed in insulin-injected FR females compared to their Adlib-insulin counterparts, indicating that over time, insulin exerts an anxiolytic effect specifically in FR females. These findings imply that in both males and females, insulin’s anxiolytic action may be dependent on prenatal adversity exposure (Fig.6g). No differences were found in entries into the light box (Fig.6h).

## Discussion

Prenatal conditions have a critical influence on fetal growth and neurodevelopment, shaping behavioral outcomes and disease risk across the lifespan ^56, 57^. These effects are partly mediated by early-life transcriptional changes in the brain, inducing a cascade of metabolic events that fuel further brain and behavioral alterations, in a self-sustaining loop ^30^, ultimately contributing to adult health consequences. Here, we show that prenatal growth restriction (FR) affects the mPFC transcriptional landscape both at baseline and in response to insulin across development, modulating emotional responsivity to environmental cues and anxiety-like behaviors. These findings enhance our understanding of the molecular mechanisms underlying the heightened emotional and behavioral sensitivity to environmental stimuli observed after prenatal adversity, such as increased psychological distress, anxiety/depression symptoms ^26–28^, and impaired value-based decision-making ^29–31^, and the modulatory role of insulin in these outcomes.

The spontaneous alternation task measures the natural tendency to alternate between cues or objects without explicit reward or punishment ^58^, a behavior that is often disrupted under high levels of stress or anxiety ^59, 60^, or by increased plasma corticosterone concentrations ^61^. We observed that FR did not affect the performance, while the known anxiolytic effect of insulin was observed in males ^16, 17^. However, in a conflict-based task (light-dark box, presenting a conflict between the drive to explore and the aversion to brightly lit areas ^55^), FR led to a sex-specific behavioral response: FR females spent less time in the lit compartment, indicative of increased anxiety-like behavior ^62^, which was completely reversed by mPFC insulin administration. In contrast, both FR and insulin infusion decreased anxiety-like behavior in males, suggesting a convergence of mechanisms. These sex-specific effects are consistent with previous findings in both FR rat models ^25, 63^ and humans born small ^31, 64, 65^, where FR influences behavior in both sexes, but through distinct pathways.

Intriguingly, in our study, insulin treatment reversed the anxiety-like behavior observed in FR females and downregulated Tmco3 expression in the mPFC. Tmco3 encodes a cation:proton transporter, classically associated with longitudinal growth ^66^. However, altered TMCO3 DNA methylation has also been implicated in several neuropsychiatric and neurodevelopmental conditions. In the human postmortem brain, changes in TMCO3 methylation have been reported in patients with Parkinson’s disease ^67^ and schizophrenia ^68^. Blood Tmco3 methylation levels have been shown to moderate the association between fetal growth discordance and cortical development ^69^, as well as diagnostic discordance for major depression in monozygotic twins ^70^, and chronic fatigue syndrome in women ^71^. Altered TMCO3 DNA methylation has also been observed in the cord blood of neonates prenatally exposed to heavy metals ^72^. Tmco3 is known to affect the startle reflex specifically in female rodents^73^ and to be upregulated in PFC microglia in a mice model of alcohol dependence ^74^, indicating that this gene influences emotional reactivity to environmental cues ^73^. Together, these findings suggest that insulin-induced downregulation of mPFC Tmco3 expression in FR females may underlie its anxiolytic effects observed specifically in this group during the light-dark test.

In FR males, insulin injection failed to induce changes in Irs2 gene expression observed in Adlib males, suggesting impaired mPFC insulin sensitivity at P90, as brain Irs2 gene expression has been shown to be upregulated by exogenous insulin ^75^. These findings are consistent with previous reports of altered brain insulin sensitivity in FR models ^24, 76^ and support the thrifty phenotype hypothesis ^8^, which posits that poor fetal growth impairs pancreatic β-cell development, leading to altered insulin secretion over the life course and increased risk for type 2 diabetes. While the brain does not rely on insulin for glucose uptake, central insulin resistance has been linked to cognitive decline, mood disorders, and neurodegenerative diseases ^21^. At P90, FR males also showed upregulation of F2rl1, which encodes protease-activated receptor-2 (PAR-2).

Given that PAR-2 blockade reduces hepatic and systemic insulin resistance ^77, 78^, its upregulation suggests the presence of central insulin resistance in FR males. Notably, PAR-2 activation has also been associated with depression-like behavior and neuroinflammation in rodents ^79^. Moreover, multiple transcription factors previously linked to insulin resistance (e.g., Glis1, Foxc2, Tbx21, Hnf4a) were enriched in the mPFC transcriptional signatures of FR animals across sexes and timepoints, corroborating the idea that the molecular response to insulin is altered in the mPFC of both male and female FR animals during the life-course.

On the other hand, it is exciting to note that insulin treatment was able to reverse some FR-induced changes in gene expression completely. In males at P90, FR induced an upregulation of Mmp8, while insulin administration led to a downregulation of Mmp8 in FR animals. Mmp8 encodes matrix metalloproteinase 8, a myeloid cell-specific proteinase that is elevated in the serum of individuals with major depression and in stress-susceptible mice following chronic social defeat; circulating MMP8 has been shown to directly infiltrate the brain, contributing to altered social behaviour ^80^. Beyond its role on major depression and stress responsiveness, a high Mmp8 expression has been associated with chronic inflammation induced by metabolic disease ^81^, especially in microglia ^82^, while its dowregulation has been linked to suppression of neuroinflammation by inhibition of microglial activation and secretion of TNF-α in the brain ^82^. Matrix metalloproteases are implicated in both pathological and adaptive brain processes: while they contribute to neuronal cell loss in neurodegenerative conditions, they also play roles in synaptic plasticity ^83^ underlying learning, memory, and various neuropsychiatric disorders (e.g., epilepsy, schizophrenia, autism, substance use disorders, and major depression particularly in response to chronic stress ^80^). In females at P21, FR induced a downregulation of Car3 (carbonic anhydrase III), while insulin administration in FR animals led to an upregulation of Car3. Car3 is involved in neuronal excitability and previously shown to be downregulated in other models of prenatal adversity ^84^. The inhibition of mPFC carbonic anhydrase activity impairs short and long-term social recognition memory in rats ^85^. Brain carbonic anhydrase activity plays a crucial role in the consolidation of extinction of emotionally salient events ^86^. The observed downregulation of this gene following fetal growth restriction, along with its upregulation after insulin administration in FR animals, aligns with the behavioral findings of reversal of heightened emotionality in FR females during the light-dark task.

In our dataset, mPFC transcriptomics associated with FR and FRINS conditions were enriched for pathways linking stress and immune responses, showing a more consolidated pattern in males at P90, while it was evident in FR females from earlier developmental time points, suggesting sex-specific differences in the timing of maturational responses to early adversity. Sex-specific differences in the timing of molecular responses are further supported by the transcriptional effects of insulin in FR animals. At P21, FR females appear more responsive to insulin, exhibiting a greater number of DEGs than FR males (Sup. Fig.2) — a scenario that reverses by P90. These sex-dependent dynamics are also evident in the broader transcriptomic patterns captured by RRHO analyses. At P0 and P21, FR males show greater divergence in insulin response compared to their saline-treated counterparts, whereas FR females display higher divergence at P90. Moreover, in FR males, there is greater concordance between the transcriptomic signatures induced by FR and those induced by insulin throughout development, where it is stronger in females at P21 (Fig.3d). Finally, sex-specific transcriptional regulation was particularly evident at P21, where genes in the Cyp family displayed the expected opposing expression patterns between males and females ^87^ (Sup. Fig.2).

Pathway analyses highlighted Wnt/β-catenin as a key pathway associated with mPFC transcriptomic changes induced by prenatal growth restriction in both males and females, and across all conditions in females. Wnt/β-catenin is a key regulator of embryonic development and adult tissue homeostasis implicated in cardiovascular and neurodegenerative diseases ^88^. (Fig.2c). Interestingly, central activation of Wnt/β-catenin signaling has been shown to elevate brain insulin levels ^89^, enhancing glucose utilization and cognitive performance in mice ^90^. Beyond the brain, Wnt signaling also modulates the susceptibility to type 2 diabetes by acting on peripheral tissues ^91^.

Chronic stress has been shown to disrupt Wnt/β-catenin signaling in the brain, leading to heightened fear and anxiety-like behaviors in mice ^92, 93^, in alignment with the behavioral and molecular alteration observed in our study.

During embryogenesis, Wnt acts as a critical morphogen guiding the development of midbrain dopaminergic neurons, including those from the substantia nigra and ventral tegmental area subtypes ^94, 95^, which project to the NAcc, striatum, and the mPFC. Wnt/β-catenin signaling, particularly in coordination with glial cells, plays an essential role in the regulation, protection, and regeneration of dopaminergic neurons ^96^. In line with this, prenatal growth restriction has been associated with disrupted development of the dopaminergic system ^97^, leading to long-lasting alterations in dopamine D1 and D2 receptor expression in the mesocorticolimbic circuit ^35, 37^, as well as reduced dopamine release in the NAcc and PFC in adulthood ^24, 35^. Here we also observed an upregulation of Drd5 (dopamine receptor D5) gene in response to insulin in adlib females at P21 and P90, supporting the well-known role of insulin modulating dopaminergic signaling ^98^, an effect that was absent in FR females (Sup. Fig.1). This aligns with evidence that Wnt/β-catenin signaling is essential for the development and maintenance of dopaminergic neurons, and further highlights it as a key pathway disrupted by FR. In humans, polygenic scores reflecting variations in dopamine signaling capacity have been shown to moderate the effects of poor fetal growth on food preferences ^99^, neurodevelopmental outcomes ^100^, and the risk of metabolic and psychiatric comorbidities during adolescence and adulthood ^101^. This convergence of evidence suggests that the Wnt/β-catenin pathway may serve as a mechanistic link between early adversity and long-term neurobehavioral and metabolic outcomes, highlighting the translational relevance of our findings.

In conclusion, prenatal growth restriction shapes behavioral and emotional responses to environmental cues, and these effects are modulated by insulin action in the mPFC in a sex-specific way. Insulin administration mimics some of the transcriptional changes induced by FR in the mPFC, aligning with our previous findings showing elevated baseline peripheral insulin levels in FR animals ^24, 25^ and supporting the hypothesis of a latent insulin resistance state following fetal growth restriction. In females, insulin reverses anxiety-like behaviours, potentially through the upregulation of Tmco3 and Car3 expression in the mPFC. The involvement of Wnt/β-catenin signaling in both the response to early adversity and insulin administration highlights this pathway as a potential mechanistic link underlying the long-term effects of fetal growth restriction. Moreover, Wnt/β-catenin signaling likely plays a key role in the altered development of dopaminergic neurons in FR animals, contributing to persistent changes in behavior and metabolic function across the life course. Given that the phenotypes observed in the FR model closely resemble those seen in humans, these findings hold strong translational potential to inform preventive and therapeutic strategies.

## Acknowledgements

This work was supported by the Natural Sciences and Engineering Research Council of Canada (NSERC, RGPIN-2018-05063 to PPS), by the Canadian Institutes of Health Research (CIHR, fellowship 193970 to PMM), by the Fonds de Recherche du Québec – Santé (FRQS, awarded to BB and PPS), by the International Collaborative Initiative in Adversity and Mental Health (AMH, awarded to PPS, PMM, and BB), and by the Canadian Neurodevelopmental Research Training Platform (CanNRT, awarded to PMM).

