## Supplemental material for "Insulin modulates mPFC gene expression and emotional behavior in a sex-specific manner following fetal growth restriction"

**Materials and methods**

**Animals.** Primiparous Sprague Dawley rats at approximately 70-80 days of age were kept in 2 per cage and maintained under standard conditions: controlled room temperature (22 ± 1 °C), 12:12 h light/dark cycle (lights on between 7:00 a.m. and 7:00 p.m.), relative humidity level between 20–50%, cage cleaning once a week, and food and water provided *ad libitum*. The estrous cycle was determined daily by vaginal smearing, and females were placed with males when receptive (proestrous). Gestation was confirmed on day 1 by visualizing the presence of sperm cells on the vaginal smear. Dams were single-housed after confirmation of gestation, and food and water were provided *ad libitum* until gestational day 10. The food restriction protocol is described below. On postnatal day (PND) 21, pups were weaned, separated into groups of two or three same-sex (same litter/group) individuals per cage, and kept in a controlled environment as previously described. Except for the cage cleaning (once a week), animals were left undisturbed from P21 until adulthood (P90).

**Food restriction model of fetal growth restriction (FR).** On gestational day 10, dams were randomly allocated into one of the following dietary groups: the control group (Adlib, n = 43), which received an *ad libitum* diet of standard laboratory chow (ENVIGO® Teklad Diets: 3.1 kcal/g, 18.6% protein, 6.2% fat, 44.2% carbohydrate (available), no sucrose), or the 50% food-restricted group (FR, n = 22), based on the model developed by ^1^. The daily amount of food provided to the FR group was determined by quantifying the mean daily intake of the Adlib group. The FR protocol occurred from day 10 of pregnancy until the pups were born. Within 24 hours of birth, all pups were individually weighed and standardized to a maximum of eight pups per litter (4 males and 4 females). Fostering was performed within this period such that all litters were adopted by Adlib dams, ensuring that the adversity experienced was limited to the prenatal period.

**Tissue collection and RNA-sequencing.** Brain tissue collection occurred at three distinct developmental stages: postnatal day 0 (P0), P21, and P90. It allowed the characterization of gene expression alterations throughout development associated with fetal growth restriction and insulin administration. On P0, the exceeding pups from both FR and Adlib litters were randomly injected with either saline (1mL/kg) or insulin (5IU/mL in saline) by intraperitoneal injection (i.p.), and had their brain collected after 30 minutes. One pup per litter was used for each group: 1 female saline, 1 female insulin, 1 male saline, and 1 male insulin. The pups were left on top of a heating pad for 30 minutes, and then sacrificed by decapitation. The brain was quickly removed, and the PFC was desiccated on ice, flash-frozen in tubes on dry ice, and stored at -80℃ for RNA-sequencing analysis. On P21, after weaning, a subsample of animals received injections with either saline or insulin (i.p.), and after 30 minutes their brains were rapidly removed, flash-frozen in isopentane, and stored at -80℃. The remaining pups were kept undisturbed from P21 to P90, when the same protocol performed with P21 pups were conducted: brain collection after insulin or saline injection. P21 and P90 frozen brains were sliced at 200μm in coronal sections using a cryostat, and brain regions were punched using a 0.8mm diameter tissue punch. In total, we have 8 experimental groups at each developmental time point (P0, P21, P90): Male Adlib Saline, Male Adlib Insulin, Male FR Saline, Male FR Insulin, Female Adlib Saline, Female Adlib Insulin, Female FR Saline, Female FR Insulin.

**RNA extraction and sequencing.** Total RNA was extracted using QIAzol lysis reagent and miRNeasy Micro kits (QIAGEN, US Cat#217084), with on-column DNase I digestion to remove residual genomic DNA. RNA was examined by Bioanalyzer 2100 (Agilent Technologies, Santa Clara, USA), and RNA libraries were prepared using Illumina TruSeq stranded total RNA LT set (Cat# RS-122-2301, Illumina Canada Ulc.). Library quality and concentration were measured using the Bioanalyzer before sequencing. Libraries were sequenced at McGill University and Genome Quebec Innovation Centre using Illumina HiSeq 4000 PE100 sequencer. Ten samples per sex, developmental time, and experimental group were sent for whole-transcriptome RNA sequencing. All RNA-seq files are available through Gene Expression Omnibus (GEO) accession GSE295966.

**RNA-seq differential gene expression analysis.** Rats RNA-Seq data were aligned to the Rattus norvegicus rn6 genome using STAR Aligner ^2^(https://github.com/alexdobin/STAR). Counts of reads aligned to rat genes were calculated in the same process by STAR with the “quantMode GeneCounts” option. Principal component analysis was used to analyze the quality and detect possible outliers in a sample. Normalization and differential gene expression analysis were performed using DESeq2 ^3^. For the initial identification of differentially expressed genes (DEGs), we applied multiple comparison corrections using a false discovery rate (FDR) threshold of < 0.05 and restricted the analysis to protein-coding genes. Volcano plots to visualize the DEGs were plotted using ggplot2, dplyr, and ggrepel R packages. To characterize broader transcriptional patterns, we applied a more inclusive threshold of uncorrected p < 0.05 and log₂(fold change) > |0.3785| (corresponding to a fold change > 30%)^4^, again limited to protein-coding genes (Venn Diagrams). Area-proportional Venn Diagrams were plotted using the BioVenn webtool ^5^.

**DEGs list comparisons.** Cell-type enrichment analysis of the final DEG lists was conducted using a rodent brain-specific gene expression atlas ^6^. A gene was classified as a cell-type-specific marker if its average expression was more than 10-fold higher than the average background expression of all other cell types within the same brain region. Enrichment was assessed by calculating the odds ratio of list overlap using the GeneOverlap package in R. Gene lists used for heatmap generation consisted of the union of all DEGs identified in any single comparison, with corresponding log₂ fold change values included even if not significant in other comparisons. Heatmaps were created using Morpheus (https://software.broadinstitute.org/morpheus), with clustering based on one minus the Pearson correlation and average linkage, and sorted by the log₂ fold change of FR-regulated genes. Enrichment analysis was performed to identify unique, similar, or common pathway maps using the 'Compare Experiments' function in MetaCore® (Clarivate; <https://portal.genego.com>). Unique sets indicate that the network object involved in a given pathway is exclusive to a specific condition. Common sets suggest that the same network object is affected across all three conditions, implicating a shared pathway component. Similar sets refer to network objects affected in two of the conditions, indicating partial overlap without being unique or fully common. Results (FDR < 0.05) were sorted by 'Differentially affected maps' and filtered by the following map categories: metabolic maps, regulatory maps, mental disorders, nutritional and metabolic diseases, and nervous system diseases. These analyses did not consider whether gene expression was up- or downregulated. Transcription factors (TFs) associated with the DEG lists were investigated using the ChIP-X Enrichment Analysis 3 (ChEA3) ^7^, ranked by the top 10 most significant. Results were visualized as bubble plots using ggplot2 and dplyr packages in R. Two-sided rank-rank hypergeometric overlap (RRHO) analysis ^8^ was performed using threshold-free differential expressed gene lists, which were ranked by degree of differential expression (DDE), where DDE represents the -log10(p-value) multiplied by the sign of the fold change from the differential gene expression analysis — DESeq2. RRHO maps were produced to evaluate concordant and discordant patterns of genome-wide overlap of the following comparisons, ran independently in each sex and timepoint: FR-saline vs. Adlib-saline compared to Adlib-insulin vs Adlib-saline and FR-insulin vs FR-saline compared to Adlib-insulin vs. Adlib-saline.

**Stereotaxic surgery to cannula implementation.** At approximately 83 days of age, the animals were anesthetized with isoflurane using an anesthesia vaporizer and placed in a stereotaxic apparatus with the incisor bar adjusted to maintain the skull horizontal between bregma and lambda. Carprofen (5 mg/kg, S.C.) was administered as an analgesic before the commencement of surgery, and 2% lidocaine (1 mg/kg) was applied locally. Ophthalmic ointment was placed on the eyes to prevent dryness during surgery. A 1 cm incision was made in the skin above the skull using a scalpel. Guide cannulas (10 mm long, 23 gauge; Plastic One, Roanoke, VA) were bilaterally lowered into the prelimbic (PL) region of the mPFC (coordinates: +3.2 mm AP, +/- 0.8 mm ML, −3.2 mm DV) for subsequent insulin or saline administration. After surgery, the animals were returned to their home cages following a 3–5-hour recovery period ^9^.

**mPFC Insulin administration.** Following a one-week recovery period after cannula placement in the mPFC, animals were bilaterally infused with a 1-μl solution of insulin (0.25 μg/μl) or a 0.9% sterile saline solution, and 5 minutes later were exposed to the first behavioural task – the spontaneous alternation task (SAT). The infusions were performed using a 1-μl Hamilton syringe attached to a polyethylene tube carrying the infusion cannula. Each infusion was conducted over a 3-minute period. To minimize peptide adhesion to the tube, the interior of the polyethylene tube was coated with rat bovine serum (Sigma). Ten days later, a second infusion occurred before the animal’s exposure to the light-dark box (LDB).

**Behavioral analysis.** Animals were habituated to the test room 30-40 minutes before the tests, and the behavioral analyses occurred in the afternoon from 1 p.m. to 5 p.m.

**Spontaneous alternation task (SAT):** At P90, animals received either insulin or saline infusions in the mPFC and, 5 minutes later, were tested in a T-maze to assess spontaneous alternation ^10^. The animals were placed in the middle of the T-maze and allowed to freely explore the apparatus for 15 minutes in a single session. The session was recorded for later evaluation using the Any-Maze software. Each arm of the T-maze was designed as A, B, and C. An arm entry was scored when all four paws of the animal were in the arm, and a spontaneous alternation (SA) was defined when the animal moved from one arm to the other two arms without repetition, i.e.: ABC, ACB, BAC, BCA, CAB, and CBA. The percentage of spontaneous alternation (% SA) was automatically calculated using the following formula: (Number of SA/(Total number of entries to the arms A, B, and C - 2)) × 100. The total number of arm entries, number of SA, and percentage of SA were evaluated throughout the 15-minute sessions in the following time points: 1, 2, 5, 10, and 15 minutes.

**Light-dark box (LDB):** At P100, the animals received a second mPFC infusion of insulin or saline 5 minutes before their presentation in the LDB to test anxiety-like behavior. The LDB is based on an approach/avoidance conflict between the drive to explore novel areas and an aversion to brightly lit, open spaces. The apparatus consists of a simple chamber divided into a dark and a light compartment connected by a small opening door. The animals were placed in the light compartment of the apparatus and were allowed to move freely between the two chambers for 15 minutes. The session was recorded for later evaluation using the Any-Maze® software, in which we stratified the analyses across time using the following minutes: 1, 2, 5, 10, and 15. We examined the duration of time the rat spent in the light and dark boxes, the number of entries in the light box, the number and duration of nose pokes from the dark into the light chamber (a measure of risk evaluation), and percentage of time doing risk evaluations ((time of pokes × 100)/time in dark).

**Cannula placement confirmation.** The day after completing the behavioral task, the animals were deeply anesthetized with a ketamine/xylazine/acepromazine mixture (0.1 ml/100 grams body weight, i.p.) and the brains were collected, stored in 10% formalin and cryoprotected in a 30% sucrose-formalin solution. Cannula placements were verified by coronal sectioning using a cryostat. Only animals with confirmed cannula placement in the mPFC were included in the behavioral analyses ^11^.

**Statistical analysis of behaviors.** A generalized linear model (GLM) was used to analyze the total 15-minute performance in the SAT and LDB, with adversity, treatment, and sex as fixed factors. To assess behavioral performance over time, we used a Linear Mixed Model (LMM) approach, which accounts for the longitudinal nature of the data and the uneven intervals between measurement time points (e.g., 60, 120, 300, 600, and 900 seconds). LMMs provide a flexible framework for modeling within-subject correlations and allow time to be treated as a continuous variable, overcoming limitations of traditional repeated measures ANOVA, such as the sphericity assumption, which is often violated with unequally spaced data. Data are presented as mean ± standard error of the mean (SEM), and statistical significance was defined as p < 0.05. All analyses were performed using SPSS Statistics version 29.0 (IBM Corp., Armonk, NY, USA), and corresponding figures were generated using GraphPad Prism version 10.2.3 (GraphPad Software, San Diego, CA, USA). We excluded animals that showed freezing behavior or that failed in exploring the light side after 1 min.

**Data availability.** RNA-seq data are publicly available in the Gene Expression Omnibus (GEO) accession GSE295966. Lists of differentially expressed genes (DEGs) are provided as source data files.

**Supplemental figure legends**

**Supplemental Figure 1: Developmental trajectories of mPFC transcriptomic signatures revealed differentially expressed genes (DEGs) that were consistently upregulated or downregulated across two developmental timepoints.** Venn diagrams depicting genes consistently upregulated or downregulated in males and females in each of the three conditions: FR **a-d**, INS **e-h**, and FRINS **i-l**. DEGs analysis considers the following comparisons: FR = saline injection in fetal growth restriction vs saline injection in adlib animals, INS = insulin injection in adlib animals vs. saline injection in adlib animals, FRINS = insulin injection in FR animals vs saline injection in FR animals. n = 9-10/group/sex.

**Supplemental Figure 2:** **Insulin injection upregulates mPFC transcription of genes related to thyroid function specifically in FR females.** Schematic timeline of experimental procedures, including the prenatal adversity model of food restriction (FR) from gestational day 10 (G10) until the end of gestation, and offspring mPFC collection for RNA-sequencing analysis in postnatal day 21 (P21) **a.** Volcano plots showing differentially expressed genes (DEGs) (FDR < 0.05) associated with FR, INS, and FRINS conditions in males **b-d** and females **h-j**. Venn diagrams represent the number of DEGs (uncorrected p < 0.05, fold change > 30%, and protein-coding genes) altered by fetal-growth restriction (FR), insulin injection in control animals (INS), or insulin injection in FR animals (FRINS). Percentage values represent the extent of overlap between conditions **e,k**. Union heatmaps display the log fold change of DEGs across matched comparisons, irrespective of statistical significance. Yellow indicates increased gene expression (positive log fold change), while blue denotes decreased expression (negative log fold change) **f,l**. Cell-type enrichment analysis of DEGs in each comparison using a curated list of cell-type-specific genes expressed in the PFC **g,m**. G = gestational day; PND = postnatal day. DEGs analysis considers the following comparisons: FR = saline injection in fetal growth restriction vs saline injection in adlib animals, INS = insulin injection in adlib animals vs. saline injection in adlib animals, FRINS = insulin injection in FR animals vs saline injection in FR animals. n = 9-10/group/sex.

**Supplemental Figure 3: Insulin injection and fetal growth restriction induce less robust transcriptional effects in the PFC of male and female offspring at birth.** Schematic timeline of experimental procedures, including the prenatal adversity model of food restriction (FR) from gestational day 10 (G10) until the end of gestation, and offspring mPFC collection for RNA-sequencing analysis immediately after birth – postnatal day (P0) **a.** Volcano plots showing differentially expressed genes (DEGs) (FDR < 0.05) associated with FR, INS, and FRINS conditions in males **b-d** and females **h-j**. Venn diagrams represent the number of DEGs (uncorrected p < 0.05, fold change > 30%, and protein-coding genes) altered by fetal-growth restriction (FR), insulin injection in control animals (INS), or insulin injection in FR animals (FRINS). Percentage values represent the extent of overlap between conditions **e,k**. Union heatmaps display the log fold change of DEGs across matched comparisons, irrespective of statistical significance. Yellow indicates increased gene expression (positive log fold change), while blue denotes decreased expression (negative log fold change) **f,l**. Cell-type enrichment analysis of DEGs in each comparison using a curated list of cell-type-specific genes expressed in the PFC **g,m**. G = gestational day; PND = postnatal day. DEGs analysis considers the following comparisons: FR = saline injection in fetal growth restriction vs saline injection in adlib animals, INS = insulin injection in adlib animals vs. saline injection in adlib animals, FRINS = insulin injection in FR animals vs saline injection in FR animals. n = 9-10/group/sex.
