## Supplementary figures and images for "Insulin modulates mPFC gene expression and emotional behavior in a sex-specific manner following fetal growth restriction"

### Supplemental figure 1

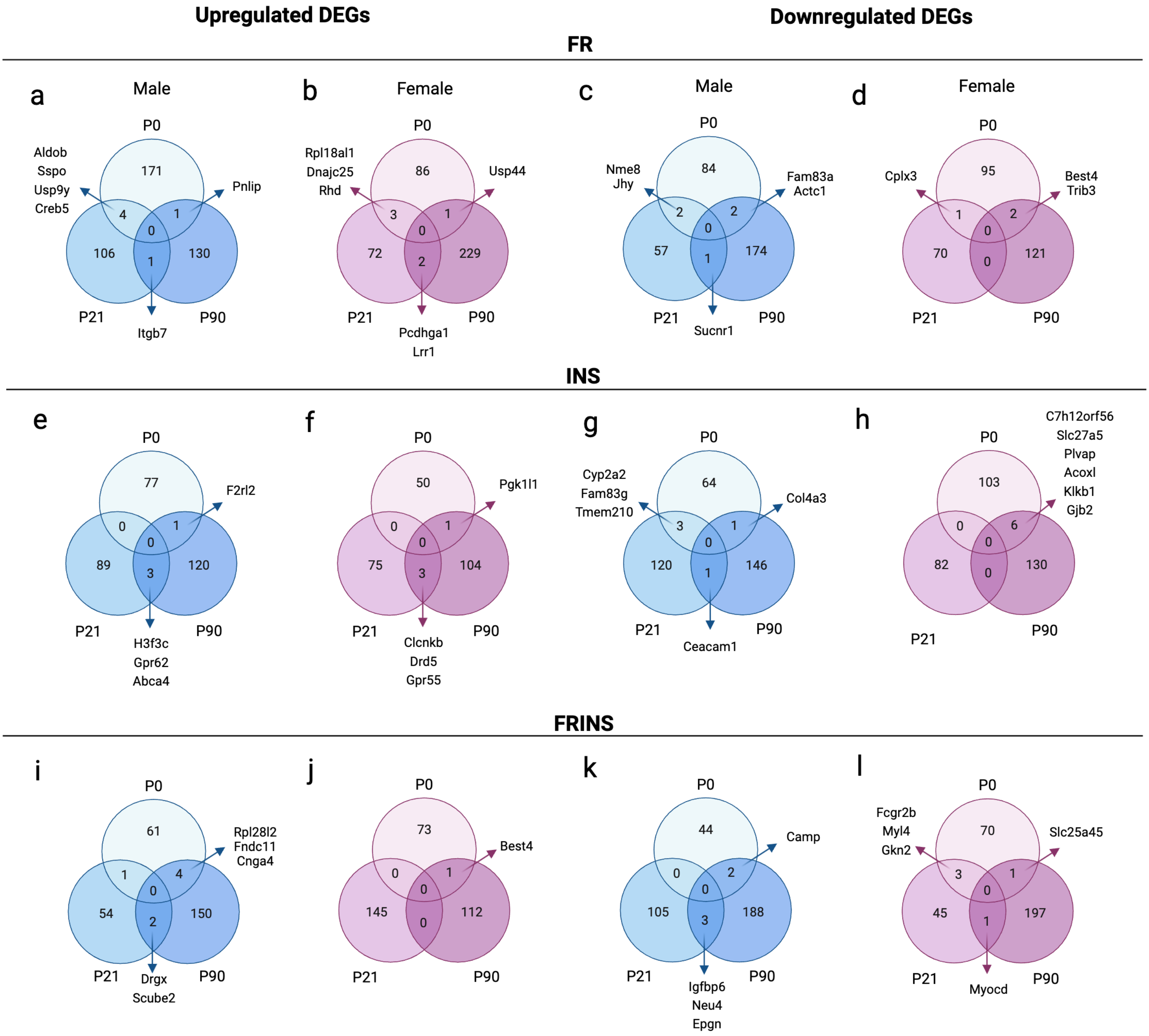

### Supplemental figure 2

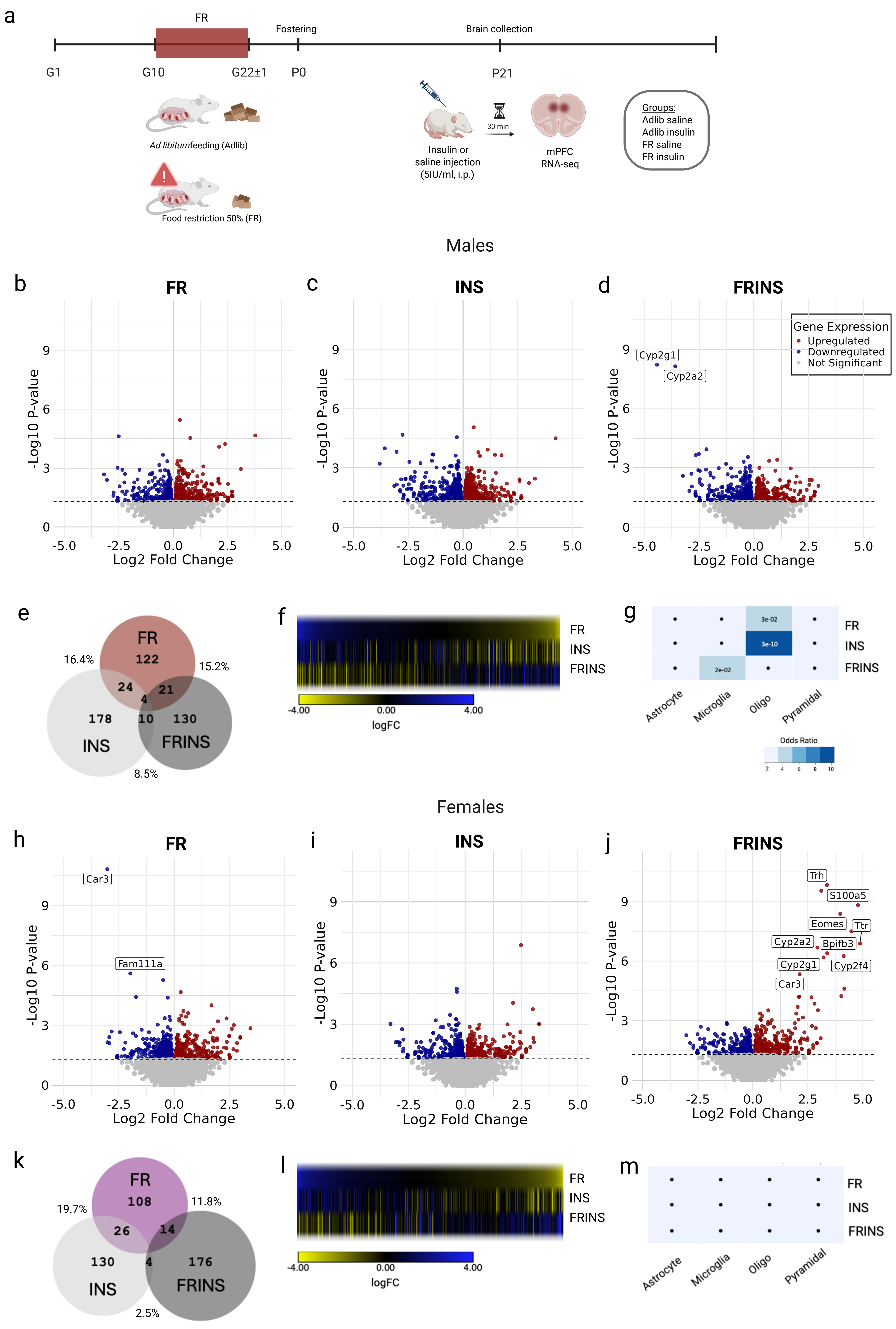

### Supplemental figure 3

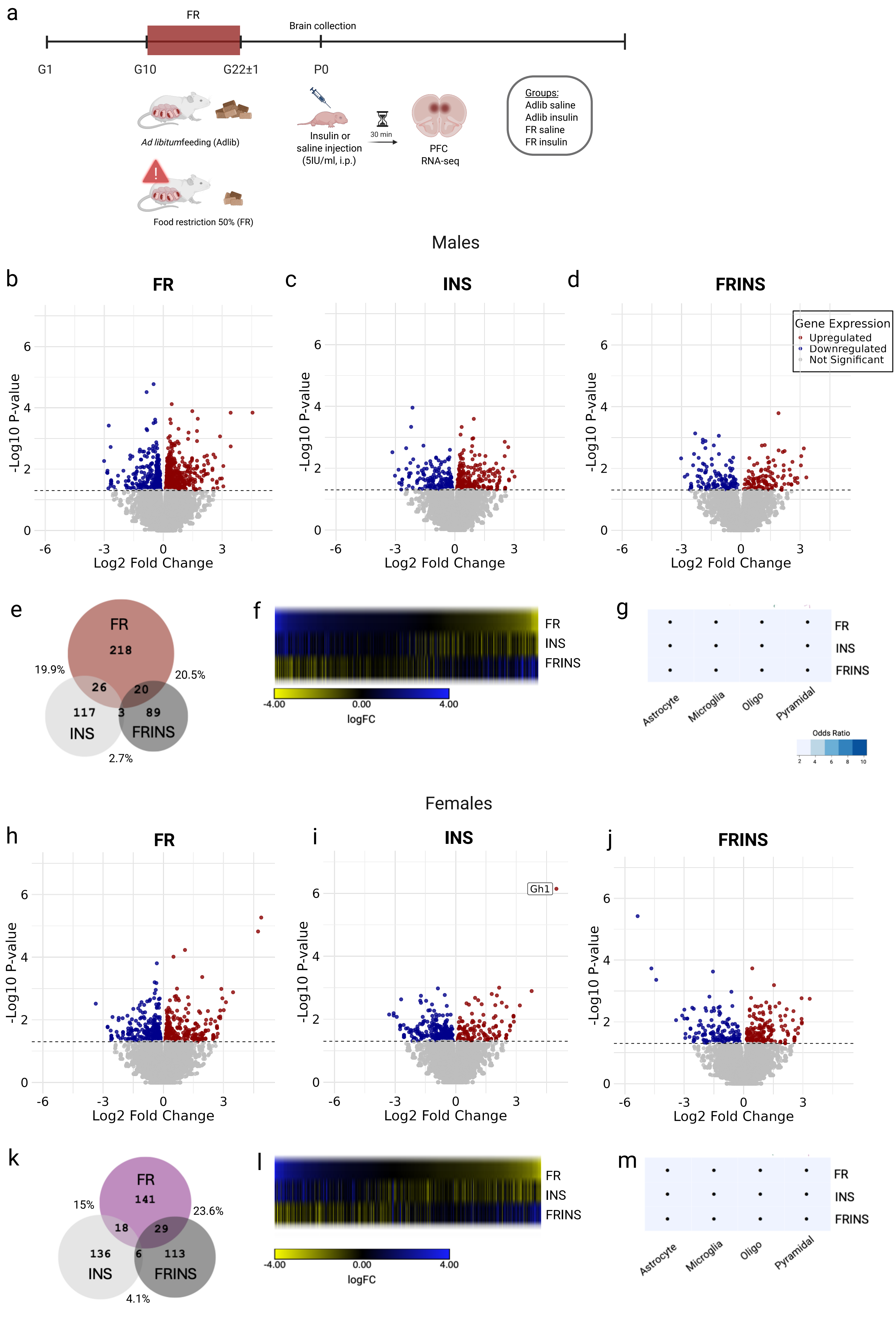
